# Mitigating Stress: Exploring how our feet change shape with size

**DOI:** 10.1101/2024.02.25.581965

**Authors:** Paige Treherne, Erin CS Lee, Michael J Rainbow, Luke A Kelly

## Abstract

If human skeletal shape increases proportionally with size (isometric scaling) this can produce exponential increases in joint contact stresses. However, if skeletal shape changes as a function of size (allometric scaling) this can mitigate increases in joint contact stress by changing the surface area to volume ratio. Here we explored whether human foot bones scale with allometry and, if so, to identify the shape features that are associated with bone size. Computed tomography scans of the two largest foot bones (talus, calcaneus) were obtained from 36 healthy individuals. We implemented a scaling analysis for each joint surface area and bone. We performed a Procrustes ANOVA to establish the shape features associated with bone size. In line with our hypothesis, articular surfaces on the talus and the posterior facet of the calcaneus all scaled with positive allometry. Interestingly, the calcaneus scaled with negative allometry, appearing more cube-like with increasing size. This may be important for mitigation of internal bone stresses with increasing skeletal size. Our findings suggest distinct, but varied scaling strategies within the foot. This may reflect the requirement to maintain healthy joint contact and internal bone stresses with increasing size.

## Introduction

Human feet exhibit a broad range of sizes and shapes (1–3). But are bigger feet just a scaled-up version of smaller feet? If this were the case, our feet (and foot bones) would scale isometrically, increasing equally in all dimensions for every unit of size increase. Isometric scaling follows the square-cube law, as length (L) increases in all directions, surface area (SA) increases to the power of two, and volume (V) increases to the power of three (SA ∝ L^2^, V ∝ L^3^) (Figure 1). However, isometric scaling is problematic for terrestrial animals because as overall size increases as a function of volume, surface area grows at a slower rate, causing disproportionate increases in joint articular contact pressures (4).

**Figure 1.**
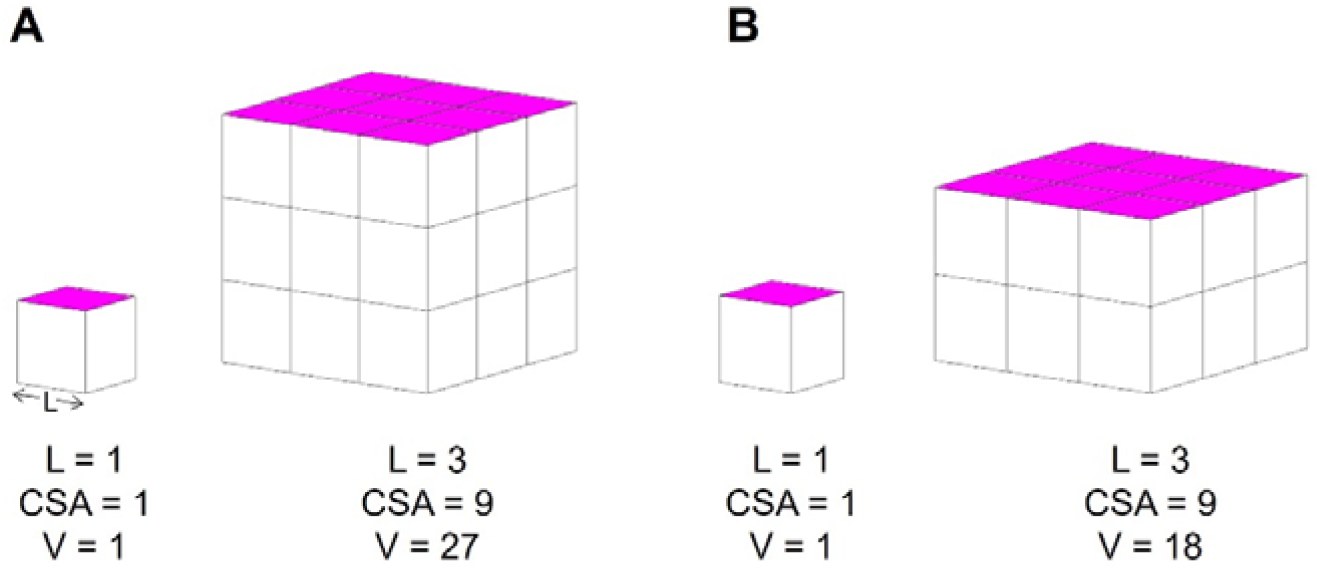
(A) Isometric scaling: the square cube law. As an object increases uniformly in length, its cross-sectional surface area (CSA, pink) increases by the length to the power of two (L^2^). The volume increases by length to the power of three (L^3^). Therefore, surface area grows at a rate of volume^2/3^, and (B) Allometric scaling: an object does not grow uniformly in all dimensions, allowing shape to change as a function of size. Following allometric scaling, surface area can increase faster than isometric scaling (positive allometry).

Allometric scaling describes a shape-function relationship that deviates from the square-cube law (Figure 1). Allometric scaling is commonly observed in terrestrial animals as a strategy to offset increased gravitational and muscular forces as animals increase in size (4,5). For example, positive allometry describes a shape function relationship whereby the surface area of a skeletal structure grows at a rate faster than would be predicted with the square-cube law for isometric scaling. This strategy has been shown to mitigate increasing joint contact stresses across a range of species (4–7). Humans exhibit a wide range of sizes; average heights of adult females around the world ranging from 150 cm to 168 cm, while the average heights of adult males range from 160 cm to 183 cm (8), and average adult body mass (combined female and male) from 58 kg to 81 kg (9). Since we still complete similar bipedal movement patterns, it is likely that we may also need to change shape as a function of size, to maintain healthy joint contact stress. This could particularly be the case for our feet, which are our primary interface with the earth we move across.

The human hindfoot complex is unique in shape compared to other non-human primates with specific adaptations to enable habitual upright locomotion (10–12). Compared to non-human primates, the human calcaneus is larger and more robust, presumably to accommodate increased ground reaction forces, and muscular forces applied by the Achilles tendon (12–14). The human talus is less ‘wedged’ with a flatter talar dome, which is hypothesized to allow for equally distributed load across the surface during locomotion with an upright tibia (15–17). Within the human population, substantial variations in hind-foot bone shape also exist (18–20). However, it is unknown to what extent these shape differences are related to size.

Therefore, the aim of this study is to explore whether human foot bones scale with allometry and, if so, identify allometry-associated shape features. First, we compared the relationships between bone volume and surface area of the calcaneus, talus, and their articular surfaces to the relationship expected from isometric scaling. To assess whether shape features are related to size we used geometric morphometrics to create a statistical shape model and regressed this model against bone and joint volume using a Procrustes ANOVA. Based on McMahon’s hypothesis that animals scale with constant stress similarity (4), we predict that the bones and their articular surfaces will scale with positive allometry (surface area growing faster than isometric scaling), and that there will be shape features at an individual bone and joint level related to size.

## Methods

### Participants

After IRB approval in accordance with the Declaration of Helsinki and informed consent, thirty-six healthy adults were included in this study (mean ± std; 171.5 ± 8.2 cm, 73.1 ± 13.7 kg; 15F). Individuals were excluded if they had a history of lower limb injury using a modified version of the Identification of Functional Ankle Instability Questionnaire (IdFAI), as well as any known neurological impairment, musculoskeletal issues, or cardiovascular conditions (21).

### Computed Tomography (CT) scans

A computed tomography scan (CT, 120 kV, 60mA, model: Lightspeed 16, n = 11, Revolution HD, n = 1, General Electric Medical Systems, USA) of each participant’s right foot was captured with the participant lying prone with their ankle in a plantar-flexed orientation (average resolution: 0.356 × 0.356 × 0.625 mm). A plantar-flexed ankle and foot orientation was chosen as it improves the in-plane resolution of the foot (22). Foot position was maintained during the scan via a custom-made foot support. Talus and calcaneus bones were segmented in Mimics 24.0 (Materialise, Leuven, Belgium) to create three-dimensional bone surface meshes (23,24). The meshes were then smoothed in Geomagic Wrap 2021 (3D Systems, SC).

### Data processing and analysis

#### Size and allometry

To test for allometry, the surface area and volume of the talus and calcaneus were computed from the tessellated meshes of each bone and articular surface (25), in Matlab (*Mathworks*, Natick, MA). To assess whether the talus and calcaneus scale with isometry or allometry, log volume (Log V) – log surface area (Log SA) plots were computed. Isometric scaling (equal growth in all dimensions) on a Log V – Log SA plot is represented by a fit line with a 0.67 slope (square-cube law, SA = V^0.67^) (4,26). Allometric scaling is found to be a deviation from either side of this line. Joint articular surface scaling was assessed using the same approach, comparing the joint articular surface area to the corresponding bone volume.

#### Joint articular surfaces

To obtain the joint articular surfaces of interest (talus sub-talar facet, talus talar dome, calcaneus posterior facet, and calcaneus anterior-medial facet), the surfaces were manually selected from the smoothed bones in Geomagic Wrap 2021 (3D Systems, SC).

#### Statistical shape models

To quantify the relationship between shape and size across our sample of tali and calcanei, we created a series of statistical shape models (Figure 2). These models were created at levels of the individual bones (talus, calcaneus) and at the level of the joint complex (sub-talar joint, combined talus and calcaneus). To initiate the alignment and point correspondence process, an initial reference bone was selected by visually inspecting all bones in the sample and choosing a reference bone that appeared to be close to an average size, with an absence of irregular features.

**Figure 2.**
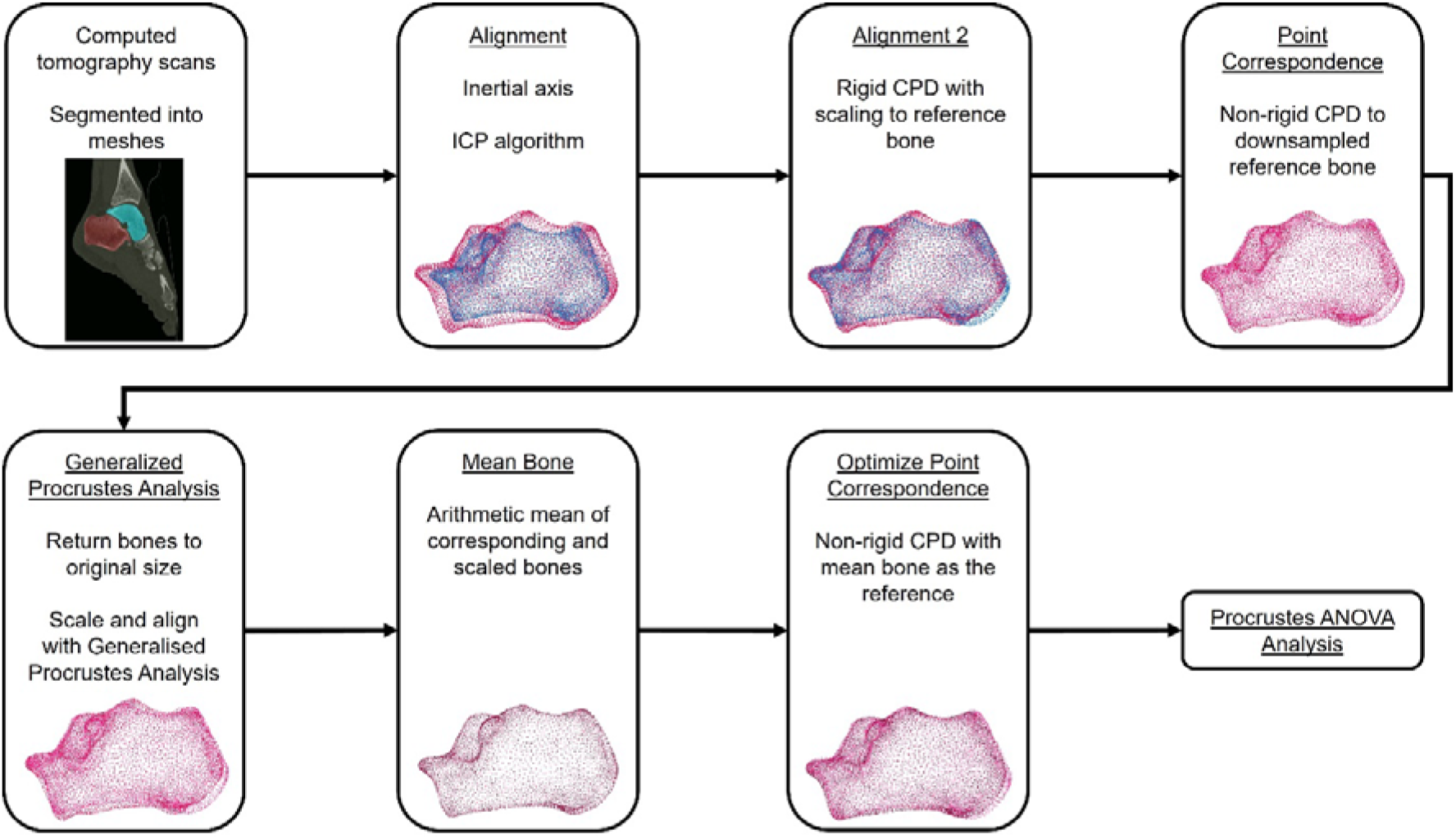
To build a statistical shape model of the sub-talar joint complex, we initially generated 3D surface meshes by segmenting CT scans. A series of alignment and point correspondence processes are undertaken to ensure robust mesh correspondence across all samples, including an iterative closest point (ICP) algorithm, and rigid and non-rigid coherent point drift (CPD) processes. We subsequently performed a Procrustes ANOVA to objectively identify bone shape features that are related to size.

#### Alignment

Initially, all bones were aligned to their inertial axes. Subsequently, bones were further aligned using an iterative closest point algorithm. A rigid coherent point drift (CPD) algorithm was then applied to align each bone in the sample to ensure optimal alignment to the reference bone while scaling the bones using relative centroid sizes (the square root of the sum of squared distances from each mesh vertex to the bone’s centroid) (27). As the meshes at this point have a different number of vertices, this scaling factor is an initial guess that will be refined in subsequent steps. These rigid transformation steps are undertaken to facilitate optimal point correspondence in the following step, without influences of alignment and size impacting the initial point correspondence.

#### Point Correspondence

Initial mesh correspondence was completed using a non-rigid CPD algorithm, reducing the mesh vertices in each bone in the sample to match the reference bone’s number of vertices, while conserving individual bone shape (27).The output bones (all with the same number of vertices) were unscaled using the scaling factor from the rigid CPD transformation to return the bones to their original size. We then performed Generalized Procrustes Analysis on the corresponding bones (28). This process scales the bones to a common centroid size and further aligns the meshes by minimizing the sum squared distances between corresponding vertices. In Procrustes space, we determined a new reference mesh by calculating the arithmetic mean of all Procrustes meshes within the dataset, for each bone (29).

Subsequently, a second non-rigid CPD was performed to re-register all bones to the mean mesh and obtain point correspondence across the sample of bones. This process yielded 3D shape coordinates, which are the set of aligned, scaled, and corresponding mesh vertices that retain only shape information. Rigid and non-rigid CPD were performed in MATLAB (27).

#### Sub-talar joint complex

The sub-talar joint complex statistical shape model was created by combining the final corresponding tali and calcanei for each participant. This allowed an analysis of the variations in shape across the entire sub-talar joint complex.

### Statistical Analysis

#### Scaling

We applied a two-tailed t-test to determine if the Log V – Log SA slope was statistically different from the isometric scaling slope (0.67). Scaling with positive allometry (slope > 0.67) means that the surface area is growing at a rate faster than that of isometric scaling, which may have implications for potentially offsetting joint pressures as humans scale. Meanwhile scaling with negative allometry (slope < 0.67) is when the surface area grows at a rate slower than that of isometric scaling.

#### Shape and size

We applied a Procrustes ANOVA to examine the relationship between size (volume) and shape for the individual bones (talus, calcaneus) and the sub-talar joint complex (talus and calcaneus combined). Procrustes ANOVA generates multivariate regression models where size is the predictor and the 3D shape coordinates are the response variables (30). A Procrustes ANOVA was applied for each individual bone (talus, calcaneus) and the entire sub-talar joint complex, using the size variables (bone/joint volume) as the predictor variables. To visualize the results of the Procrustes ANOVA, the mean shape was warped to the fitted values (shape coordinates) predicted by each regression model across the spectrum of the size variable used, e.g., smallest volume to largest volume, highlighting the regions of each bone that change shape with increasing or decreasing bone size. Procrustes ANOVA analysis was performed with the R packages *geomorph and RRPP* (31–34).

## Results

### Scaling Results

The surface area and volume of the talus scaled isometrically (slope Log V – Log SA = 0.67, p = 0.34, Figure 3A) while the calcaneus scaled with negative allometry (slope Log V – Log SA = 0.62, p = 5.56e-36, Figure 3B), meaning that as bone volume increased the surface area grew more slowly than would be expected from isometric scaling.

**Figure 3.**
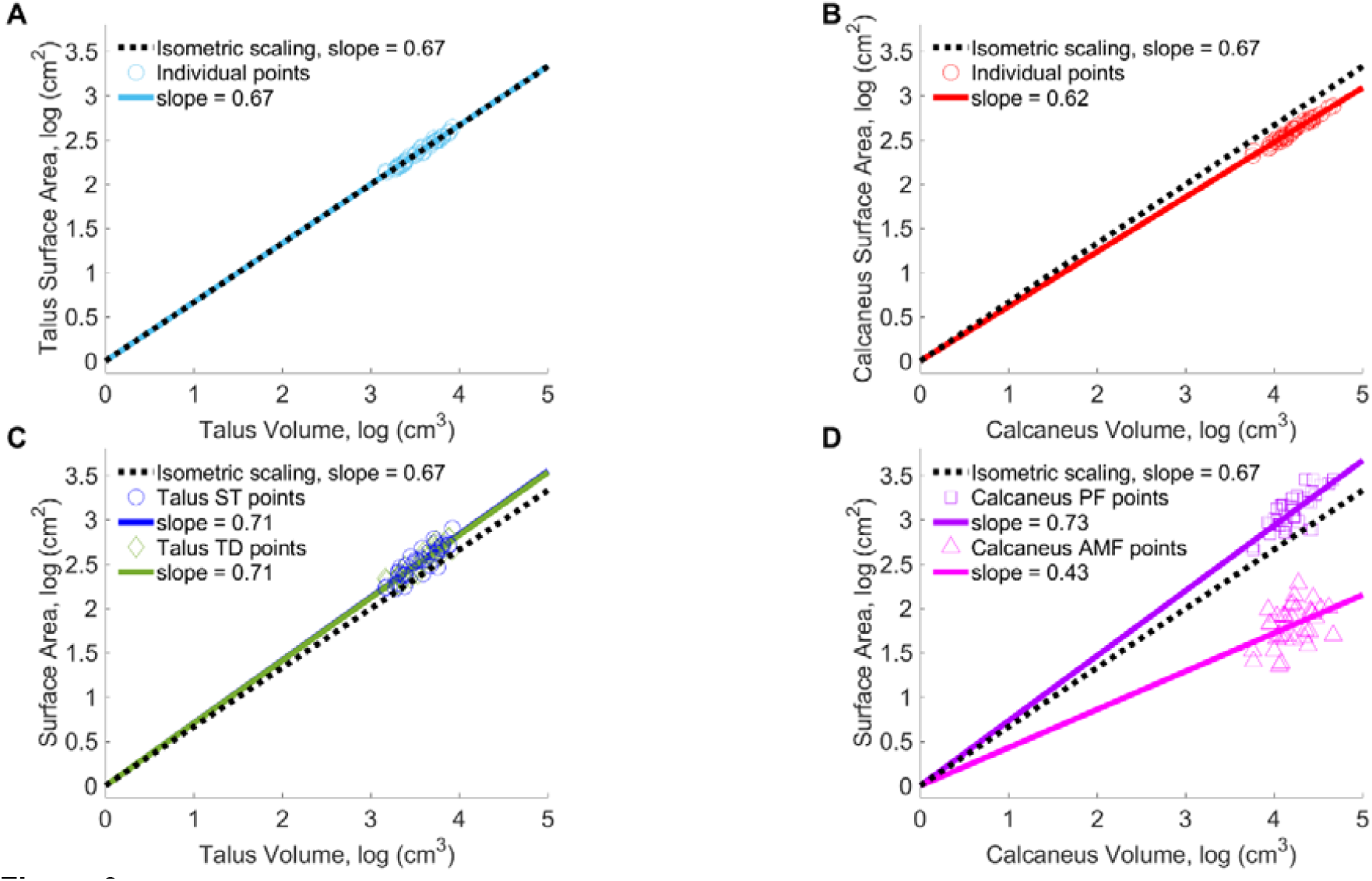
Scaling plot (Log V – Log SA) of (A) the talus with individual participant points and line of best fit in blue, (B) the calcaneus with individual participant points and line of best fit in red, (C) talus articular surface scaling plot, with sub-talar (ST) surface individual participant points and line of best fit in light blue, talar dome (TD) individual participant points and line of best fit in green, and (D) calcaneus articular surface scaling plot, with the posterior facet (PF) individual participant points and line of best fit in purple, anterior-medial facet (AMF) individual participant points and line of best fit in pink. The isometric scaling line (slope = 0.67) is in black across all graphs.

The articular surfaces on the talus (Figure 4A) both scaled with positive allometry, with the surface area scaling increasing at a greater rate than would be expected with isometric scaling (slope Log V – Log SA = 0.71 of both the sub-talar facet and talar dome, sub-talar facet p = 4.38e-13, talar dome p = 6.55e-16, Figure 3C). The posterior sub-talar facet on the calcaneus (Figure 4B) also scaled with positive allometry (slope Log V – Log SA = 0.73, p = 5.67e-15, Figure 3D). Contrastingly, the anterior-medial facet on the calcaneus scaled with negative allometry (slope Log V – Log SA = 0.43, p = 1.38e-24, Figure 3D).

**Figure 4.**
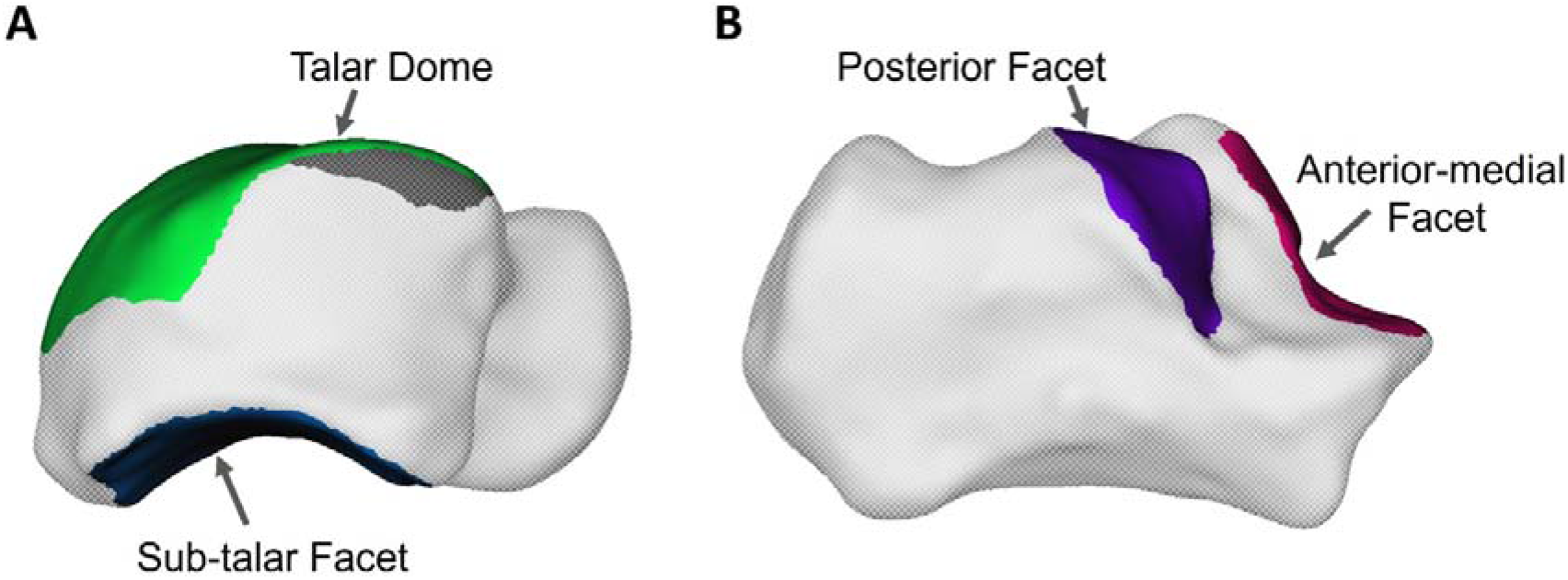
Articular surfaces of (A) the talus, talar dome in green and sub-talar facet in blue, and (B) calcaneus, posterior facet in purple and anterior-medial facet in pink.

### Relationship between bone shape and bone size

We observed no relationship between talus shape and volume (size) (p = 0.07, R^2^ = 0.04, Z = 1.5). The shape features of the calcaneus were related to calcaneus volume (p = 0.003, R^2^ = 0.06, Z = 2.7). As the calcaneus increased in size, the bone appeared to become relatively taller (superior-inferiorly) wider (medio-laterally), and shorter (antero-posteriorly). Collectively, these shape features appear to make the calcaneus become more ‘cube-like’ as it becomes larger (Figure 5). In addition, the inferior aspect of the cuboid facet is elongated while becoming more curved.

**Figure 5.**
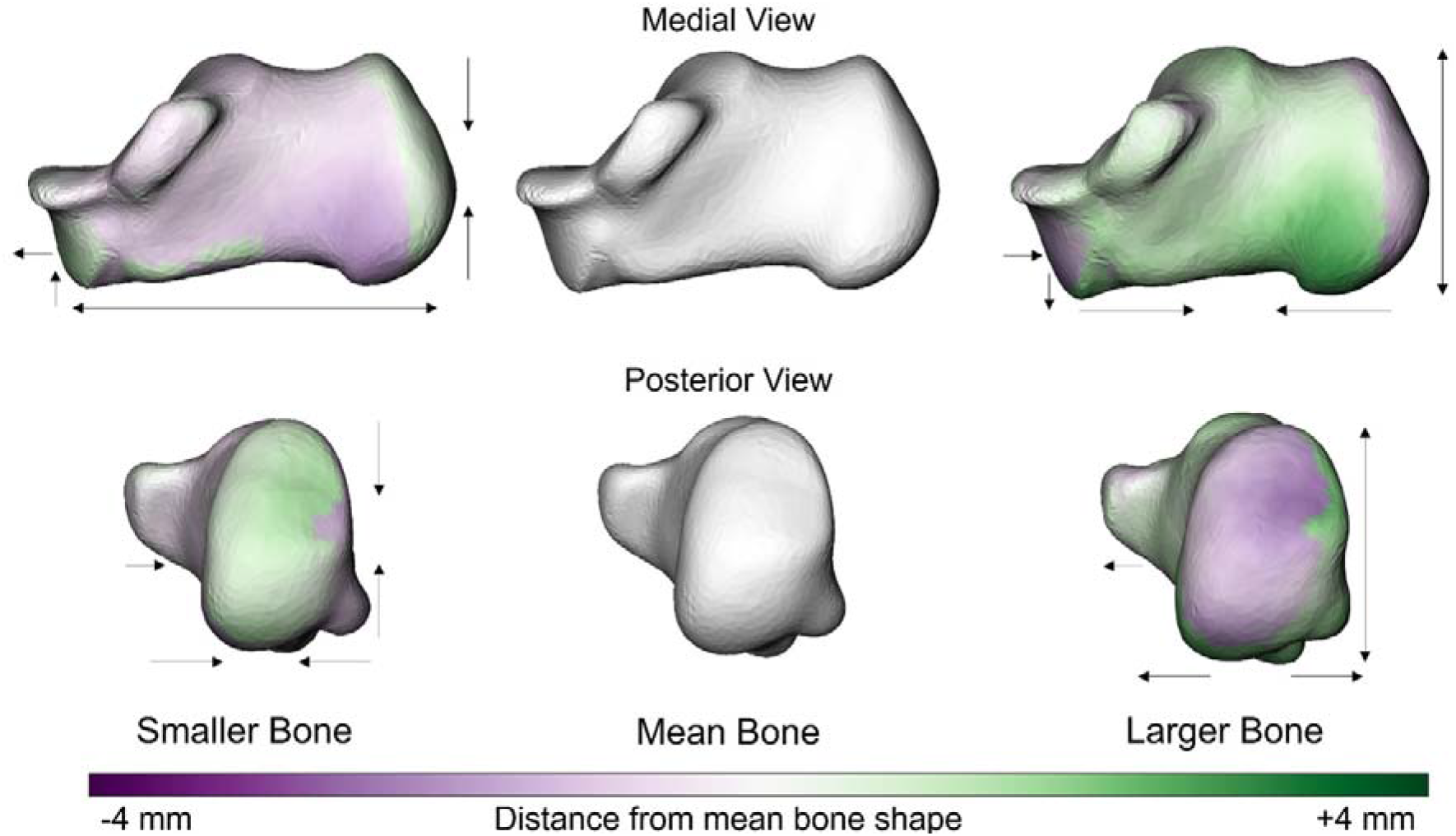
Anatomical shape variations of the calcaneus linked to calcaneal volume. Green indicates where the bone is larger and outside the mean shape, while purple shows where the bone sits inside the mean bone. The arrows focus on the location and direction of change (pointing at the bone indicates decreased size of a feature, whereas an arrow pointing away from the bone indicates increased size of a feature).

Shape features of the sub-talar joint complex were also related to size (combined calcaneus and talus volume, p = 0.001, R^2^ = 0.06, Z = 3.1). As the joint-complex becomes larger we qualitatively observe similar shape changes in the calcaneus to those observed at the individual bone level. The calcaneus became relatively taller, wider, and shorter with increasing joint volume. (Figure 6). Additionally, the joint-level analysis indicated shape-size relationships at the talus, with a relative shortening of the posterior process and the talar dome growing in width less than what would be expected for isometric scaling, appearing to become narrower as the joint-complex increases in size.

**Figure 6.**
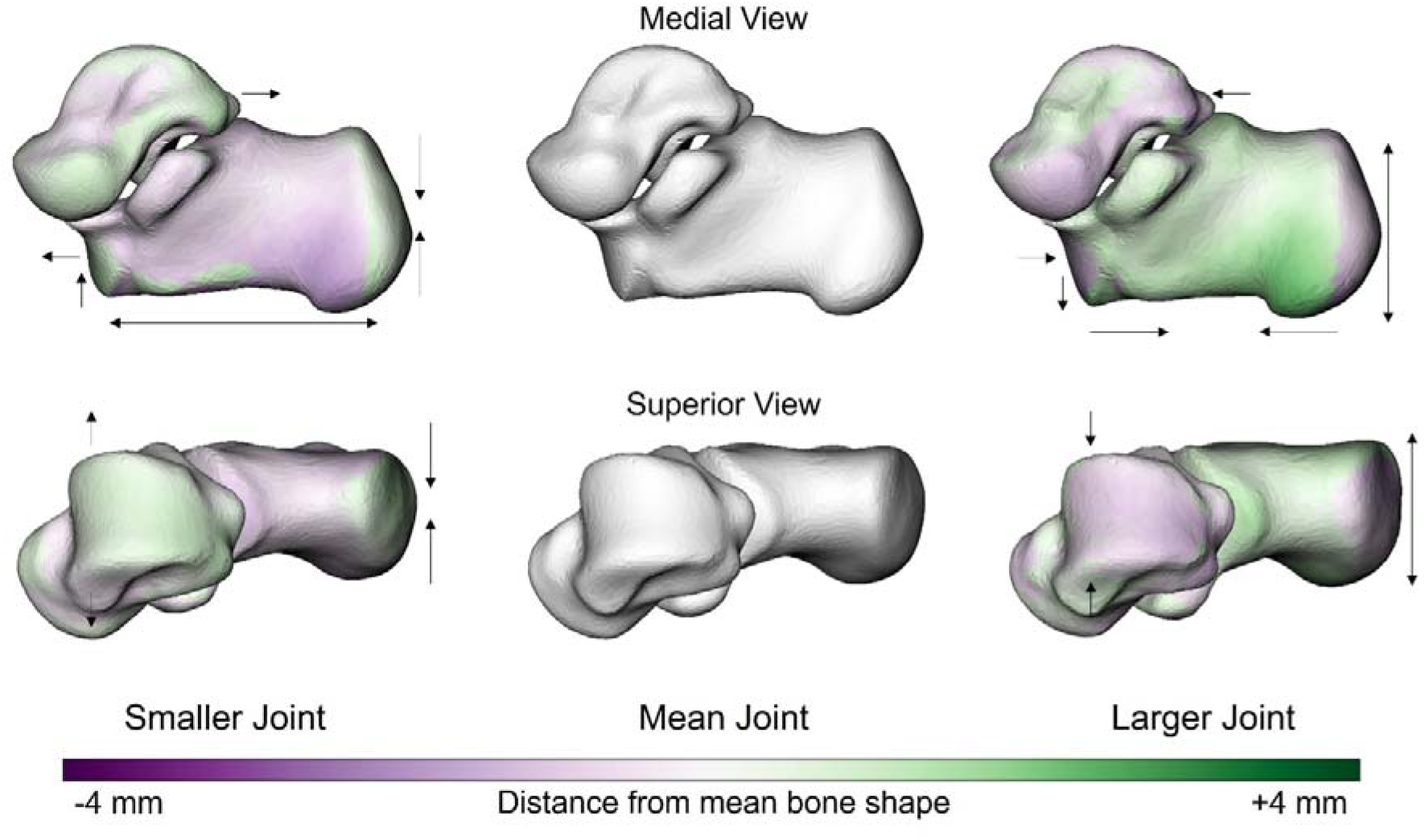
Anatomical shape variations of the sub-talar joint complex linked to sub-talar joint volume (combined calcaneus and talus volume). Green indicates where the bone is larger and outside the mean shape, while purple shows where the bone sits inside the mean bone. The arrows focus on the location and direction of change (pointing at the bone indicates decreased size of a feature, whereas an arrow pointing away from the bone indicates increased size of a feature).

## Discussion

Human tarsal bones vary widely in shape and size. We sought to understand if the tarsal bones change shape as a function of size, to inform our understanding of ankle and sub-talar joint contact mechanics during locomotion. As hypothesized, we found distinct shape-size relationships at the levels of the articular surface, individual bones, and the sub-talar joint complex. However, the nature of these relationships varied, with differing scaling patterns and shape-size relationships occurring across each structural level. Despite this variability, it appears that shape features of the calcaneus dominate the shape-size adaptations of the sub-talar joint complex.

The allometric variations for most of the articular surfaces deviated from the scaling behavior of the bones themselves. We found that the talus isometrically scales; however, the talar dome and sub-talar facet (talus) scaled with positive allometry, indicating that the surface area of the joint grows at a relatively faster rate than would be expected under isometric conditions. The calcaneus and the anterior-medial facet scaled with negative allometry, while the posterior facet on the calcaneus scaled with pronounced positive allometry. These findings are largely consistent with the theory that the joint surface area needs to grow at a faster rate to maintain relatively constant joint contact stresses across a range of skeletal sizes (4).

Clear shape-size relationships were observed for the calcaneus, but not for the talus. Given that the talus scaled isometrically, this result is unsurprising. Interestingly, as the calcaneus grows in size, it becomes taller (supero-inferiorly), wider (medio-laterally), and relatively shorter (antero-posteriorly). The medial calcaneal tubercle (the origin of the plantar fascia) becomes enlarged (Figure 5). Collectively, these shape features appear to transition the calcaneus to a more cube-like (robust) shape as it increases in size. This observation aligns with our finding of negative allometric scaling in this bone, as becoming more cube-like would lead to a relatively lower growth in surface area as would be expected under isometric conditions (4).The calcaneus is encumbered with large magnitudes of repetitive forces applied from the ground due to gravity, and also from muscular forces (e.g. Achilles tendon) (35). Given that gravitational and muscular forces scale with size (4,5), a more cube-like shape may improve the capacity of the bone to contend with increasing internal bone stresses (4,7).

The calcaneus and talus do not function in isolation, but rather as an articular complex (sub-talar joint). Therefore, we sought to understand how these two bones change shape together as a function of size. Unlike the individual bone analysis, we observed specific shape features in both the talus and calcaneus that were related to size. The shape features observed in the calcaneus as part of the sub-talar joint complex were similar to the features observed at the individual bone level. The observed shortening of the talar posterior process and widening of the talar dome appear to be in response to the relative increase in width and decrease in length of the calcaneus with increasing size (Figure 6). Given that the shape features of the calcaneus are similar at a bone and sub-talar joint complex level, it appears that the shape changes in this bone are driving the shape features across the entire complex. The talus sits within the ankle joint mortise and has no direct muscular attachments or direct interaction with the ground. Therefore, being constrained by the surrounding bones (calcaneus, tibia, navicular) may provide less opportunity for the talus to change shape without concomitant changes in the other surrounding bones.

Some methodological limitations should be considered when contextualizing our findings. We have not included sex as a covariate in our statistical model due to sample size. We chose to quantify size using bone volume and joint volume (combined talus and calcaneus volume) because foot size and body height do not have a strictly linear relationship. However, this may limit the ability to infer the relationship between our observed shape changes and loads associated with body size. Future studies that incorporate function and direct or indirect measures of joint contact mechanics may improve our understanding of the shape, size and function relationships within the human foot.

In summary, we observed articular surface, bone, and joint complex level shape-size relationships. These relationships are variable and appear to reflect the competing demands of balancing healthy joint contact mechanics and internal bone stress. These findings have important implications for the development and progression of degenerative joint conditions such as osteoarthritis. Further research exploring how joint posture and mobility interact with shape and size to determine joint contact mechanics have the potential to enhance management of these conditions via surgical and conservative means.

